# Healthy microbiome - moving towards functional interpretation

**DOI:** 10.1101/2023.12.04.569909

**Authors:** Kinga Zielińska, Klas I. Udekwu, Witold Rudnicki, Alina Frolova, Paweł P Łabaj

## Abstract

Microbiome-based disease prediction has significant potential as an early, non-invasive marker of multiple health conditions linked to dysbiosis of the human gut microbiota, thanks in part to decreasing sequencing and analysis costs. Microbiome health indices and other computational tools currently proposed in the field often are based on a microbiome’s species richness and are completely reliant on taxonomic classification. A resurgent interest in a metabolism-centric, ecological approach has led to an increased understanding of microbiome metabolic and phenotypic complexity revealing substantial restrictions of taxonomy-reliant approaches. In this study, we introduce a new metagenomic health index developed as an answer to recent developments in microbiome definitions, in an effort to distinguish between healthy and unhealthy microbiomes, here in focus, inflammatory bowel disease (IBD). The novelty of our approach is a shift from a traditional Linnean phylogenetic classification towards a more holistic consideration of the metabolic functional potential underlining ecological interactions between species. Based on well-explored data cohorts, we compare our method and its performance with the most comprehensive indices to date, the taxonomy-based Gut Microbiome Health Index (`GMHI`), and the high dimensional principal component analysis (`hiPCA)`methods, as well as to the standard taxon-, and function-based Shannon entropy scoring. After demonstrating better performance on the initially targeted IBD cohorts, in comparison with other methods, we retrain our index on an additional 27 datasets obtained from different clinical conditions and validate our index’s ability to distinguish between healthy and disease states using a variety of complementary benchmarking approaches. Finally, we demonstrate its superiority over the `GMHI` and the `hiPCA` on a longitudinal COVID-19 cohort and highlight the distinct robustness of our method to sequencing depth. Overall, we emphasize the potential of this metagenomic approach and advocate a shift towards functional approaches in order to better understand and assess microbiome health as well as provide directions for future index enhancements. Our method, **q2-predict-dysbiosis (Q2PD)**, is freely available (https://github.com/Kizielins/q2-predict-dysbiosis).

## Introduction

The prevalence of a range of diseases and conditions peripherally or directly linked to microbiome health such as Inflammatory Bowel Disease (IBD), diabetes, obesity and even various cancers, continue to increase globally and substantial funds are currently spent on diagnosis and treatment (M’koma, 2013, Hong et al., 2019). While a correlation between gut microbiome composition and human health is widely acknowledged (Vijay and Valdes, 2022), the accurate identification of microbial and host markers of disease states remains elusive. Accordingly, the ability to evaluate patient health status based on a gut microbiome snapshot would be of high clinical value. Stool-based methods are promising because they can be collected non-invasively and frequently, and analysis time is short. Furthermore, decreasing costs of stool-sample analysis via next generation sequencing makes such microbiome characterization a strong competitor as a diagnostic tool (Bajaj et al., 2020).

Dysbiosis, defined as a perturbation of gut homeostasis, is believed to be accompanied by reduced microbiota diversity and increased prevalence of ‘harmful’ bacteria in adults (DeGruttola et al., 2016, Hrncir, 2022). Eubiosis (opposite of dysbiosis) can be perturbed by a wide range of factors including infection, diet, exercise, antibiotics, stress or poor sleep (Martinez et al., 2021). The simplest interventions currently applied for the prevention or alleviation of mild microbiome dysbioses include dietary modification or prebiotics (often non-digestible food types which promote the growth of beneficial microorganisms), ingested live bacteria or probiotics (beneficial bacteria usually in capsules), and lifestyle changes. More severe cases of gut dysbiosis, failing to respond to the above interventions, may qualify for fecal microbiota transplants, FMTs, which are increasingly gaining traction in clinics worldwide (Bull and Plummer, 2015). However, host-microbiota and intra-microbiota interactions are both extremely complex and highly individual, and despite success with FMT’s, we have as yet no real understanding of why or how they work.

There are a number of accepted approaches used for the evaluation of a given gut microbiome’s health status based on stool composition. Alpha diversity (Shannon entropy, for example) is a frequent choice, as microbiome richness was long believed to be a key driver of microbiome health and robustness (Li et al., 2022, Gong et al., 2016). Beta diversity has also been applied in a number of longitudinal studies, albeit to a lesser degree and mainly to identify eubiotic samples based on a time-resolved proximity to other healthy samples (Zouiouich et al., 2021). The most robust index to date, outperforming diversity indices, is the Gut Microbiome Health Index, or the `GMHI` (Gupta et al., 2020), recently renamed the Gut Microbiome Wellness Index (`GMWI`). An updated version of this index has recently been published (Chang et al., 2024). The GMHI is based on the ratio of 50 microbial species associated with healthy or unhealthy gut ecosystems and is reported to exceed 73% accuracy in determining disease state, thus the authors suggesting that gut taxonomic signatures can predict health status. Another metagenomic gut-health index expanding on the `GMHI` approach, `hiPCA` (Zhu et al., 2023), was introduced as a monitoring framework for personalized health purposes. The personalized approach is achieved by analyzing the contribution of each bacterium to the index, which allows for the identification of high-influence (ostensibly keystone) species in different patient groups. The `hiPCA` claim of better performance than the `GMHI` is attributed to the authors’ application of additional transformation and clustering algorithms. Importantly, such studies are often defined by datasets limited in scope to industrialized nations and thus a less than complete consideration of diet-environment-microbiome interactions.

However, a recently re-visited definition of the microbiome emphasizes the importance of not just the microbiota (a community of microorganisms), but the whole “Theatre of Activity”, ToA (Berg et al., 2020). This ToA includes structural elements (proteins, lipids, polysaccharides), metabolites and environmental conditions (Lee et al., 2024). It is tightly bound to its corresponding ecological niche, and the synergistic relations between species provide all the necessary, community defining components. Based on this definition, we maintain that an index constructed from taxonomy alone is hardly sufficient to accurately capture biological phenomena occurring within the gut environment – the key to understanding gut dysbiosis. Instead, we hypothesize that **to effectively determine health:** i) a metagenomic functional profile is required (microbiome phenotype); and ii) species interactions (e.g measured as co-occurrence) but not just presence should be considered.

We introduce an approach that is based on identifiable metagenomic features within ecosystems that extend beyond diversity measures and basic taxonomic information. This **function-**centricism is broached in two ways: i) directly by evaluating the functional potential within and between species and ii) indirectly by assessing co-occurrence and synergism between bacterial species. Our goal is not only to distinguish between healthy and diseased, but importantly, to also quantify the degree of dysbiosis in each sample for the given cohort. We derive the health-describing features based on an exploratory analysis of healthy samples from the Human Microbiome Project 2 (HMP2, Lloyd-Price et al., 2019) and outperform **Shannon entropy**, the `GMHI` and the `hiPCA` in healthy versus inflammatory bowel disease (IBD) and obese classification. The robustness of our index is further validated by corroboratively, classifying two additional IBD-focused cohorts. By retraining the IBD-specific parameters on an additional set of 30 diverse cohorts encompassing a range of diseases, we demonstrate the superior performance of our approach. Our findings reveal that function-, rather than taxonomy-based features are more informative for the accurate classification of biological samples. Additionally, our method effectively identifies longitudinal microbiome changes in COVID-19 patients, which the `GMHI` and the `hiPCA` are unable to capture, and, crucially, it is <u>distinctly robust</u> to sequencing depth. Our method `q2-predict-dysbiosis (Q2PD)` is freely available (https://github.com/Kizielins/q2-predict-dysbiosis).

## Results

### High prevalence of “core functions” in health

In order to develop a strategy to assess the degree of dysbiosis in a given microbiome sample, we must define eubiosis, i.e. the healthy microbiome. We based our initial analysis (described in the Methods section) on 384 medically-determined healthy samples from the HMP2 project and identified the most prevalent species, regardless of abundance (Figure 1a). We observed that 50% of species were present in less than 5% of samples, and hardly any species were shared by all individuals. On the other hand, the prevalence of functions within the healthy population had an opposite trend - 50% of functions were already represented by at least 40% of individuals (Figure 1b), a functional redundancy unaccounted for in the `GMHI` (or the `hiPCA` which is based on it). These results convinced us further about the unsuitability of basing an index purely on the presence of “core taxa” and encouraged a shift of focus towards more prevalent functions instead.

**Figure 1.**
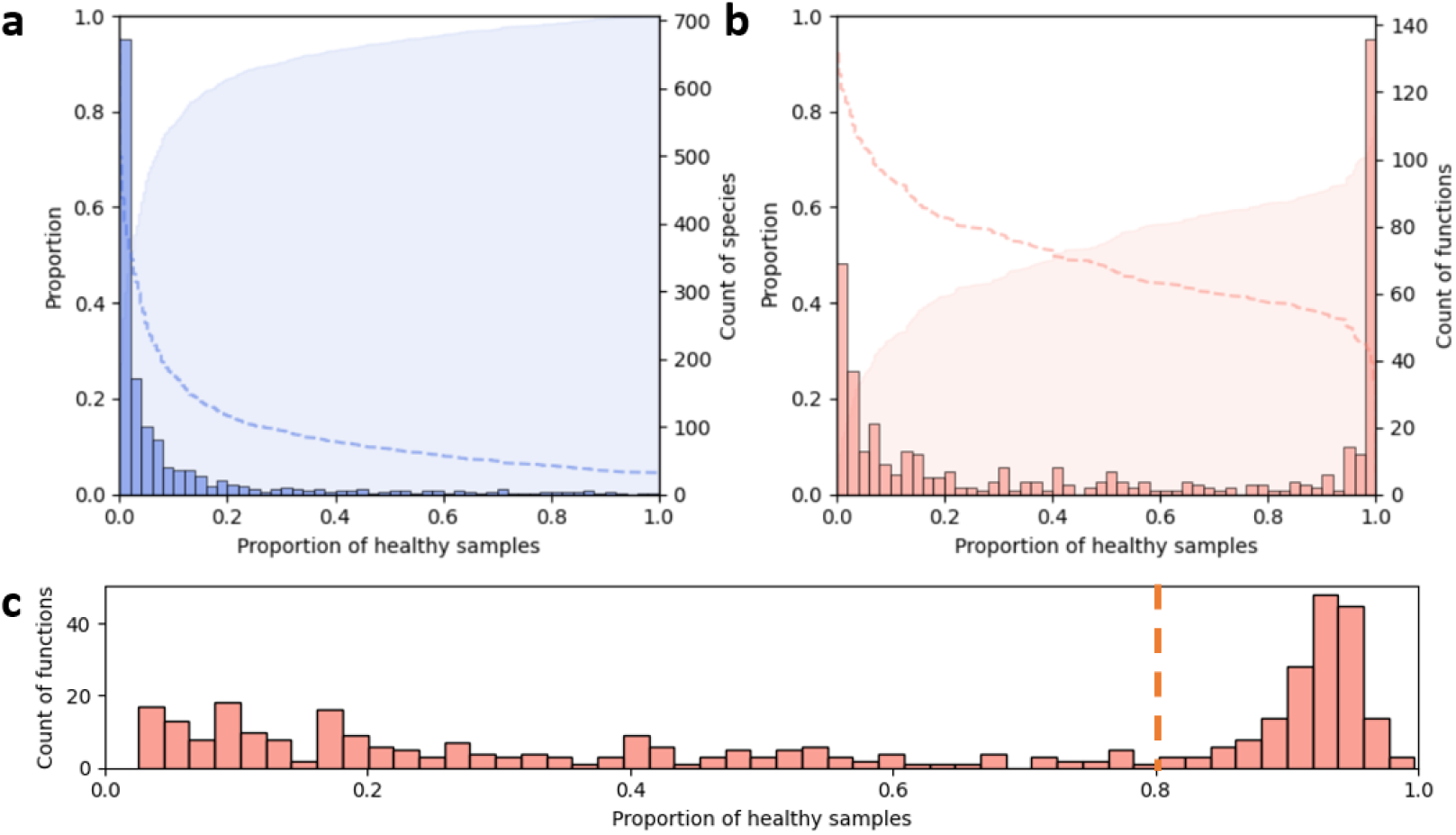
Distributions of species **(a)** and functions **(b)** present in healthy samples from the HMP2; absolute values of species and function counts are shown as histograms (with scales on the right-hand side, with shaded cumulative sum in the background and an inverse of the cumulative sum represented with a dashed line (with scales on the left-hand side). **(c)** Distribution of functions in healthy individuals from the HMP2 and two validation cohorts.

The addition of healthy samples from two validation cohorts maintained the function distribution profile obtained solely with HMP2 samples (Figure 1c). In order to test whether the functions were universal or cohort-specific, we calculated separately the distributions for functions present in 1, 2, or all 3 cohorts. We found that all functions missing from at least 1 cohort were present in less than 10% of samples, which indicated the presence of high-prevalence functions in all three cohorts. Based on the increased occurrence of certain functions in over 80% of samples (dotted line in Figure 1c), we defined them as “core functions” (refer to Supplementary Table 1 for full list). According to MetaCyc classification, 73.5% of “core functions” were assigned as “Biosynthesis” pathways, 18.8% “Degradation/Utilization/Assimilation”, and 7.6% “Generation of Precursor Metabolites and Energy”. In addition, a few of the above were additionally classified as “Superpathways”. The classification aligned well with a previously reported high prevalence of carbohydrate and amino acid metabolism-related pathways, potentially forming the functional microbiome core (Zou et al., 2019). A detailed analysis of the core functions identified in our study, however, is out of the scope of this manuscript.

Shannon entropy calculated on species and functions allowed for good discrimination between healthy and unhealthy samples in the HMP2 dataset (Figure 2a top). However, the trends were unclear in the two validation cohorts (Figure 2a middle and bottom), reflecting the need for more complex methodology, expanding beyond microbiome richness, in order to classify datasets without obvious separation. Diversity analyses revealed that the number of functions per sample remained similar, or even increased, during microbiome transitions from healthy state to dysbiosis in HMP2 (Supplementary Figure 1). While admittedly, this cannot be measured solely using metagenomics data, the similarity could hypothetically be due to altered expression of genes usually silenced in the eubiotic state, although this observation was not reproduced in the validation cohorts. Next, we investigated the presence of “core functions” in different groups, testing whether “core functions” are maintained or replaced by others in dysbiotic, IBD, samples. While the differences were not significant in most cases, we noted a visibly higher percentage of “core functions” and a higher percentage of all “core functions” in healthy as compared to disease samples (Supplementary Figure We carried out differential enrichment analysis using LEfSe (Segata et al., 2011), performed separately for each cohort, and identified 100 functions that were more abundant in healthy as compared to disease cohorts (Figure 2b, Supplementary Figure 2). Over 90% of functions enriched in healthy samples were “core functions”, while they constituted less than 5% of functions enriched in the unhealthy class of validation cohort 2 and HMP2 (Figure 2c). The LEfSe analysis on validation cohort 1 revealed only 10 significantly enriched functions (7 in health and 3 in disease), all of which were core. This indicated a more heterogeneous functional landscape within this cohort.

**Figure 2.**
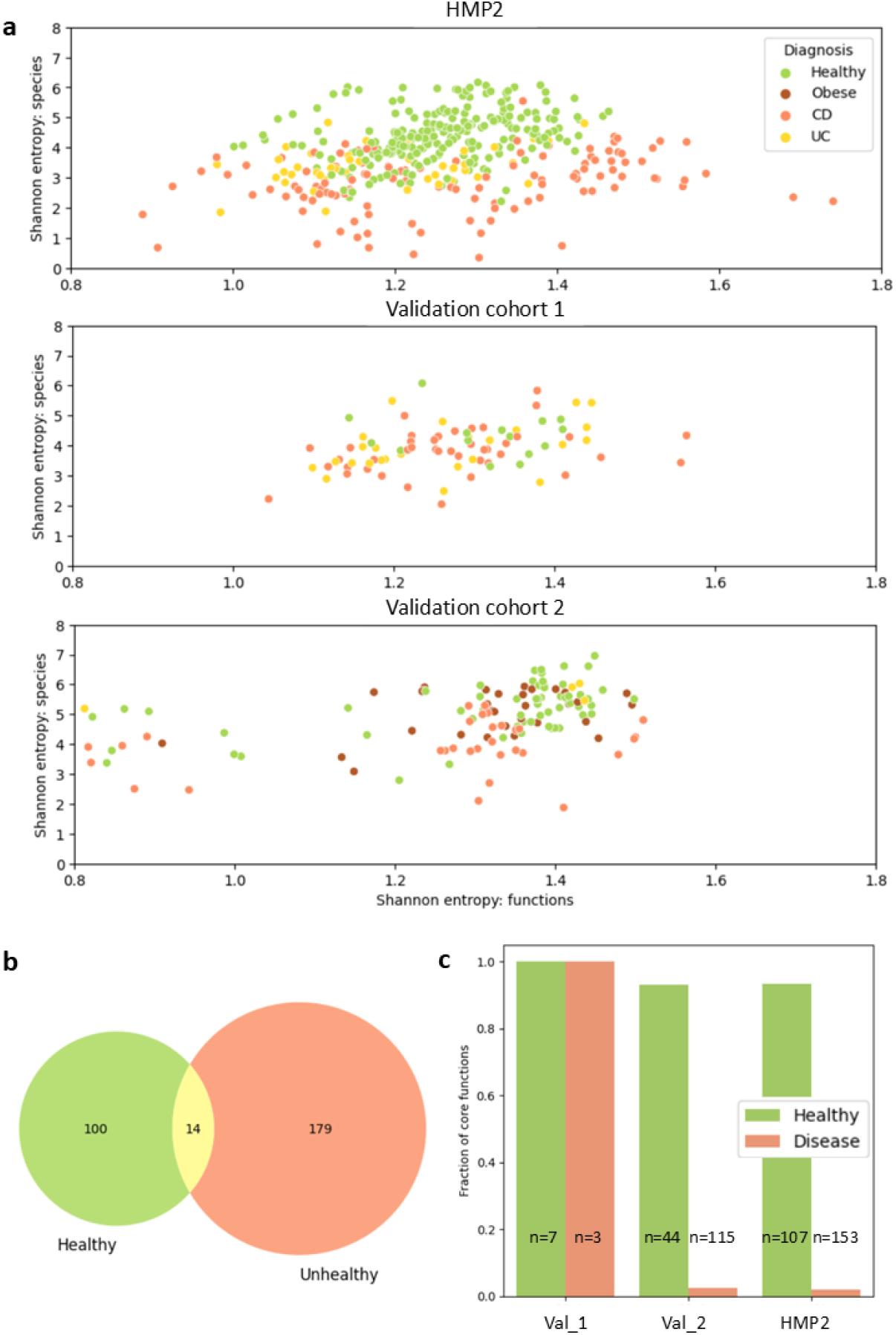
**(a)** Shannon entropy scores for species and functions in healthy and unhealthy samples from the HMP2 and validation cohorts. **(b)** LEfSe differential enrichment analysis: overlap of enriched pathways in healthy and unhealthy individuals in the validation and HMP2 projects. **(c)** Fraction of core functions among differentially enriched functions in healthy and unhealthy individuals in the validation and HMP2 cohorts.

### Species interactions and function contributions in health

Corroborating results from past studies, we observed a decrease in the species abundance in dysbiotic samples (Supplementary Figure 3, Mirsepasi-Lauridsen et al., 2018, Mosca et al., 2016). Having previously noted an increase in the number of functions (Supplementary Figure 1), we speculated that the remaining species may contribute to core or new functions, forming new connections with one another. Due to the substantial number of initial connections to analyze (170 “core functions” and 1490 species present in at least two projects), we restricted the number of species to those most informative in the context of health-/ disease-state separation. We chose the Multi-Dimensional Feature Selection (MDFS) algorithm, as it was the only feature selection method accounting for inter-feature interactions that we were aware of at the time of manuscript submission (Piliszek et al., 2019, Mnich & Rudnicki, 2020). This approach reduced the number of relevant species to 587 allowing us to eliminate noise and focus on the most important interactions (see Methods for more details about the feature selection procedure).

We then used the SparCC algorithm, designed specifically for compositional data, to investigate correlations between the MDFS-selected species in health and disease (Friedman and Alm, 2012). We did not observe any trends in the number of correlations, or in the fraction of positive correlations per group, that would indicate differences between the two. However, we identified opposite relationships of some species in different groups (Figure 3a). A number of species we found to be positively correlated in eubiosis and are generally considered beneficial (e.g. *Eubacterium rectale, Faecalibacterium prausnitzii*, and a number of *Bacteroides species*), and those relationships would be disrupted in dysbiotic groups. We observed that the prevalence of the pairs positively correlated with health was higher than in a number of disease-associated groups (Figure 3b). Due to this, we included the co-occurrence of such species as another feature of interest to aid potentially in the determination of microbiome health.

**Figure 3.**
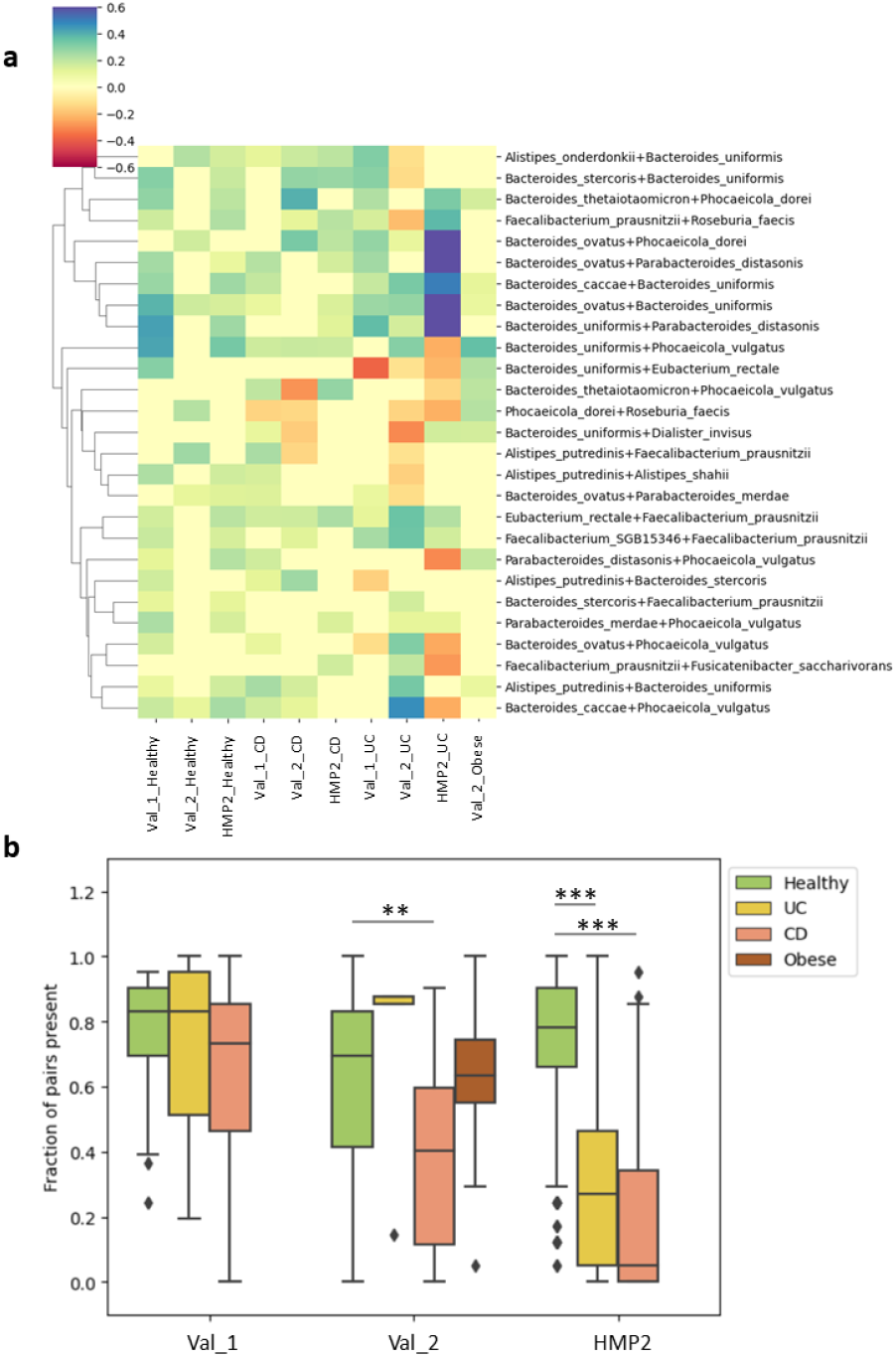
**(a)** `SparCC` correlation strengths between species, restricted to pairs which were not negatively correlated in any healthy cohort. **(b)** Prevalence of the pairs in different cohorts.

Based on our previous results, we hypothesized that the contributions of each species to functions would be relatively stable in the healthy state and less predictable in disease. To test this, we compared the contributions of MDFS-identified species to “core functions” in different groups (Supplementary Figure 4). We did not observe any differences between health and disease, despite a relatively tight clustering of the healthy groups. However, we found stronger results when exploring functional redundancy. While the average number of species per function and the average number of functions per species did not reliably separate healthy from diseased profiles (Supplementary Figure 5), the latter approach was more informative as described in detail below. This finding was congruent with our earlier suspicions of an inherent functional plasticity of microbiome structure; modulation of function altering connectivity in the interaction network, leading to a shift towards less abundant, non-core functions upon perturbation of homeostasis. It also highlighted the challenge of identifying dysbiosis based on singular features, which were never statistically significant for all cohorts – and the utility of multiple perspectives for microbiome description, in order to meaningfully classify them.

### Testing the accuracy of prediction for healthy and IBD individuals

Our final set of health-defining microbiome features included the following parameters (details of how each feature was calculated can be found in Table 1):

**Table 1.** Parameters of the Q2PD model.

| Name | Explanation | Derivation |
| --- | --- | --- |
| Frac_of_core_functions_found | fraction of "core functions" found | number of "core functions" in a sample / number of all "core functions" in our list |
| Frac_of_core_functions_among_all | fraction of "core functions" among all functions | number of "core functions" in a sample / number of all functions in a sample |
| Species_found_together | fraction of species pairs commonly occurring together in healthy samples | SparCC correlation of species > 0.1, only nonnegative correlations in healthy cohorts |
| Func_contributions_per_species | average number of function contributions per species | number of all species to functions contributions based on stratified output / number of all species in a sample |
| GMHI_good | number of "good" GMHI species | list of health-associated GMHI species |
| GMHI_bad | number of "bad" GMHI species | list of disease-associated GMHI species |

i. the fraction of “core functions” found,
ii. the proportion of “core functions” among all functions,
iii. the proportion of co-occurrent species pairs in healthy samples,
iv. the average number of functional ‘contributions’ per species.

In addition, we included two parameters derived from the GMHI method – the number of “good” and “bad” GMHI species identified in a sample, which would enable us to compare between approaches. We then fed these described parameters into a machine learning model. We opted for a Random Forest classification algorithm (Breiman, 2001) due to its robustness to imbalanced data and interpretability, and performed a leave-one-out cross-validation. For the model training, validation and subsequent testing, we used taxonomic and functional profiles from the curated Metagenomics database (Pasolli et al., 2017) to ensure consistency and reproducibility (see Methods).

Our index demonstrated the strongest statistically significant separation between healthy and Crohn’s disease or ulcerative colitis individuals across all methods evaluated (Figure 4a). Alongside hiPCA, it was one of only two approaches to achieve a statistically significant distinction in the Nielsen_2014 cohort. Overall, both our index and hiPCA exhibited comparable levels of accuracy and AUC across the three cohorts, outperforming the other methods by a notable margin (Figure 4b). By contrast, the GMHI consistently delivered mediocre results, and Shannon entropy only performed well applied to the HMP2 cohort, yielding low AUC scores (0.37–0.53) in all other cases. When index values were used as features in the Boruta algorithm (Kursa & Rudnicki, 2010), with health status (0/1) as the target variable, our index emerged with the highest mean and summed importance across all cohorts (Figure 4c). The above highlights the ability of our index to provide the most informative scores for health status prediction as compared to the other methods.

**Figure 4.**
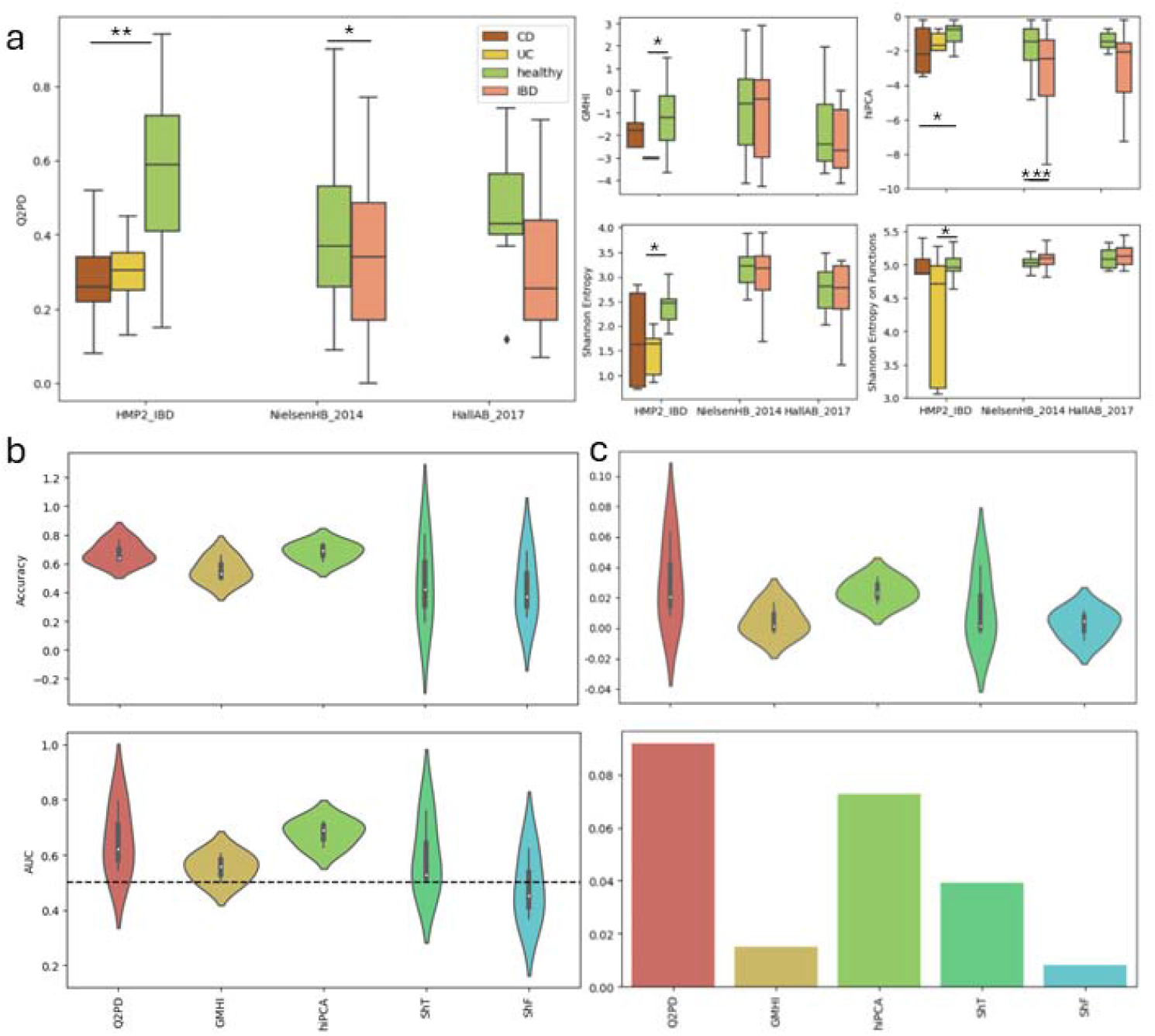
**(a)** `Q2PD`, `GMHI, hiPCA` and `Shannon entropy (on species and functions)` scores for healthy and IBD (Crohn’s disease = CD, ulcerative colitis = UC) individuals. **(b)** Accuracy and AUC values for each index, per IBD cohort. ShT = Shannon entropy on taxa, ShF = Shannon entropy on functions. **(c)** Average (top) and summed (bottom) importance of each index in the context of IBD prediction, determined by Boruta.

### Beyond IBD

While only healthy and IBD individuals had erstwhile been included in the development and validation of our approach, we wondered about the applicability of the `Q2PD` to dysbiosis attributed to other diseases. We extended our dataset to another 27 additional cohorts from various disease states, equating to 30 datasets used for method validation. We did not change any parameters of the **Q2PD**, which were originally determined based on the healthy samples from the HMP2. Instead, we retrained the model with the new data and performed a leave-one-cohort-out approach to ensure a robust benchmark. The procedure placed our index at a disadvantaged position, as some of the added datasets had been used to develop and train the other methods and our approach was thus truly blind to outcome in these new cases.

Despite the disability, and gratifyingly, the `Q2PD` achieved in terms of performance, the highest average accuracy and AUC across all datasets {(AUC=0.61, accuracy=0.58) > hiPCA (AUC=0.58, accuracy=0.57) > GMHI (AUC=0.55, accuracy=0.55) > Shannon entropy on species (AUC=0.52, accuracy=0.53) > Shannon entropy on functions (AUC=0.44, accuracy=0.43)} (See Figure 5a). The consistently poor performance of both entropy-based measures suggested their highly limited utility as predictive indices.

**Figure 5.**
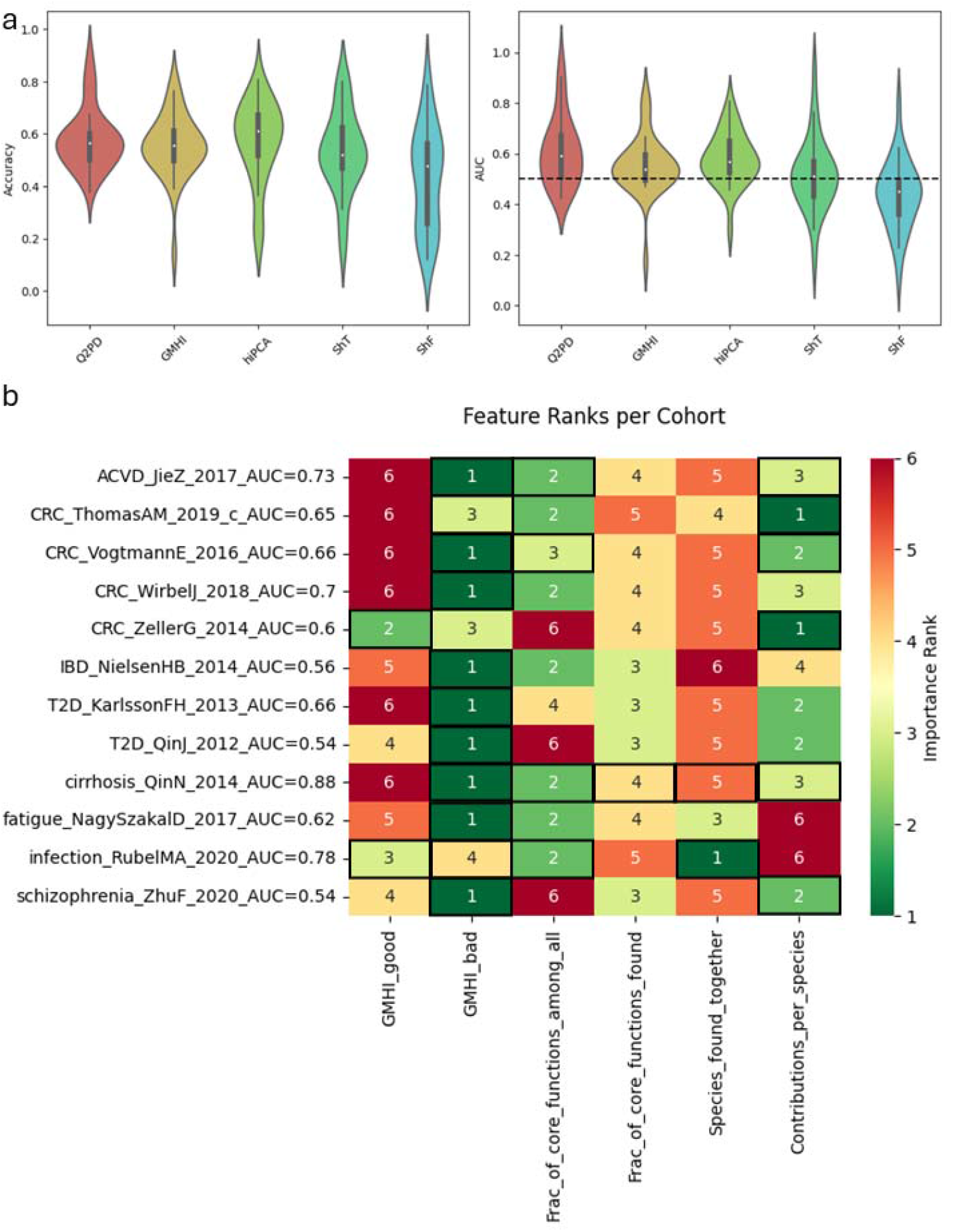
**(a)** Average accuracy and AUC values for each cohort. **(b)** Average importance ranks of features for each cohort. A lower rank indicates greater importance. Cells with black borders indicate variables identified as informative by the Boruta algorithm. The AUCs were produced as a part of the feature importances analysis when training on individual cohorts, and therefore do not align with the AUCs produced by the Q2PD. Only datasets with AUCs > 0.5 are shown.

We observed that where `Q2PD` classified a particular cohort better than the other indices, it did so with a significantly greater margin than when it lost to the other methods. Its average winning AUC margin over the hiPCA, the second best classifier, was 0.19 while the losing margin to the hiPCA was 0.09 when the hiPCA had the highest AUC. The average winning AUC margin of Q2PD against the mean of the other indices was 0.20 and 0.11 if any other of them was better. In both cases, the T-test statistics for the differences between the means of the Q2PD’s winning AUC margins and that of the hiPCA or the average of others produced p-values of 0.03 and 0.02, respectively, indicating a significant classification improvement with our method in areas in which the remaining indices did not classify well. The improvement was even more striking when we excluded datasets that had been used for the training of either method. In this case, the winning margin of the Q2PD was 0.22 while the losing margin to the mean of the other indices if any of them was better was 0.09.

Our investigation into the accuracy and AUC of the indices for each cohort revealed substantial variability in terms of the classes of cohort that each index was able to classify (Supplementary Figure 6). We observed that while some cohorts such as Liss_2016 could be classified well with function-based indices (Q2PD and Shannon entropy on functions), other cohorts such as Gupta_2019 were slightly better separated with taxonomy-based indices (hiPCA, GMHI and Shannon entropy on species). An exploration of the importance of feature(s) associated with the model training on each dataset alone revealed a large amount of diversity, suggesting different kinds of information used to classify different cohorts (Figure 5b). Interestingly, when trained on the level of individual cohorts, a random forest would pick the “GMHI_bad” as its most informative parameter (scoring the lowest rank in 9 cases) and the “GMHI_good” as the worst (appearing at the bottom of the ranking 6 times). The importance of the function-based features would vary depending on the dataset, with “Contributions_per_species” winning and losing twice and the other two being consistently in the middle of the ranking.

Having observed a discrepancy between the superior Q2PD performance and the greatest importance of the “GMHI_bad” parameter, we investigated this further by testing which types of diseases were best suited for each index to utilize. To this end, we designed a ranking method based on the number of times each index achieved the highest AUC for the most cohorts for each disease. Overall, Q2PD outperformed the other indices for 6 diseases (atherosclerotic cardiovascular disease, colon cancer, infection, metabolic disease, schizophrenia and fecal microbiota transplant: donor versus patient classification). This was followed by hiPCA for 5 (Behcet’s disease, IBD, type 2 diabetes, chronic fatigue and cirrhosis), Shannon entropy on taxa for 2 (Parkinson’s disease and acute diarrhoea) and GMHI for 1 (a cohort with mixed diseases). From this we also calculated an average rank for each method for all the diseases above and yet again, Q2PD ranked best (average ranking=2.39), followed by hiPCA (2.71), Shannon entropy on taxa (3.14), GMHI (3.32) and Shannon entropy on functions (3.57). The poor performance of the GMHI indicated a diminished role of the “GMHI_bad” parameter when combining all datasets, and implied a better generalization of the health status using function-based parameters.

### Q2PD robustness to longitudinal alterations and sequencing depth

To construct a final model, we trained the random forest classifier on the complete set of 30 cohorts. We then took advantage of a longitudinal COVID-19 dataset that had been sequenced both shallowly and deeply, and was ‘unseen’ by any methods, in order to evaluate performance of Q2PD across sequencing depths. The set consisted of three groups – COVID-19 patients who, during the course of the treatment, were either i) transferred to the intensive care unit (ICU) or ii) recovered (noICU), and iii) controls (healthy hospital staff). For every individual, two timepoints were selected - “1” that was collected upon hospital admission, or in the case of staff, early in the pandemic, and “2” which was the final sample taken from each individual. As expected, sequencing data with fewer than 300,000 reads per sample fell short in accurately classifying individuals, primarily because the limited read depth did not provide sufficient coverage for comprehensive functional annotation (Figure 6a).

**Figure 6.**
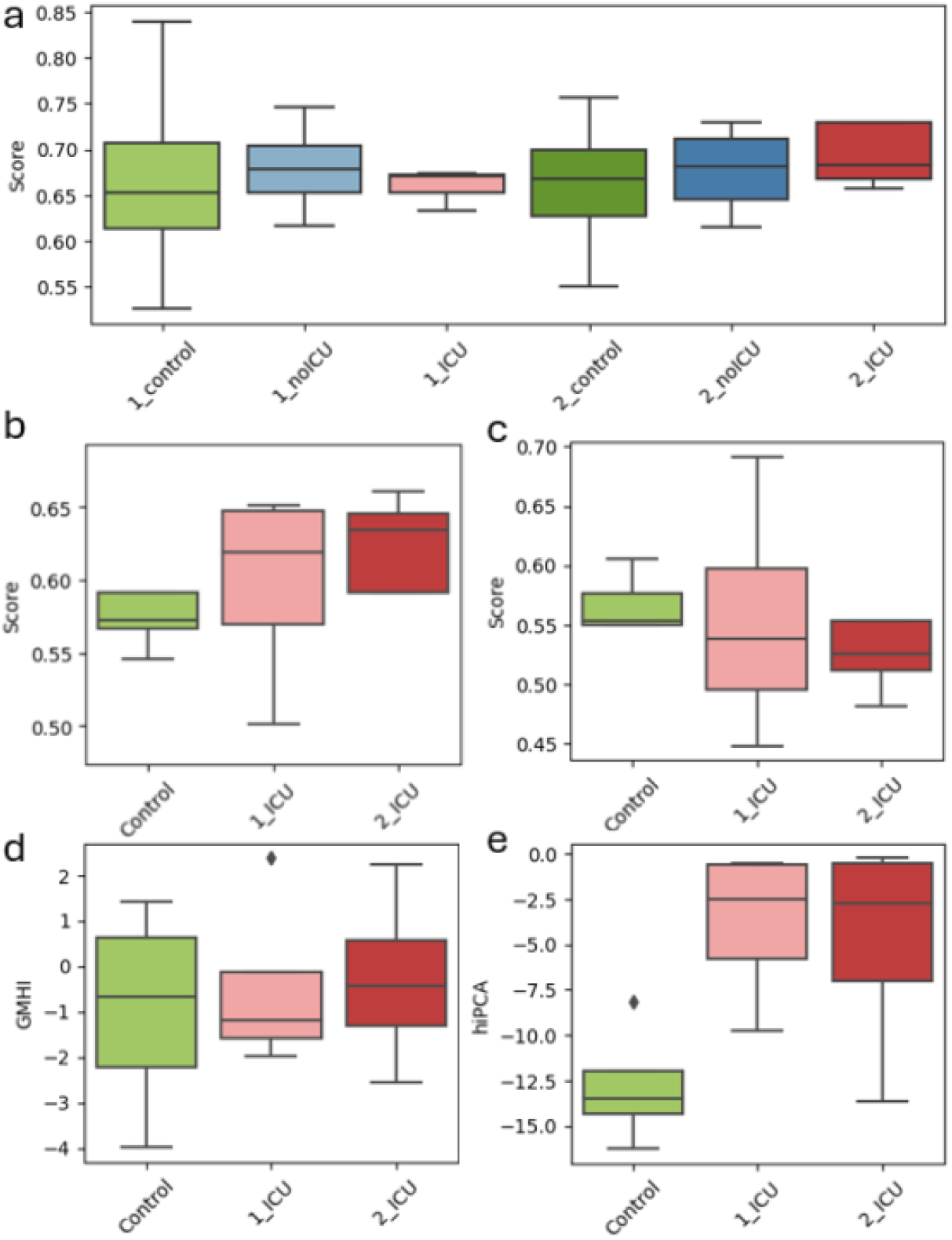
**(a)** `Q2PD` predictions on the shallow COVID-19 cohort. **(b)**. `Q2PD` predictions on the deep COVID-19 cohort. **(c)** `Q2PD` predictions on the deep COVID-19 cohort with the taxonomic parameters excluded. GMHI **(d)** and hiPCA **(e)** predictions on the deep COVID-19 cohort.

Surprisingly, despite its earlier described successes at classification, Q2PD was not able to distinguish between healthy and COVID-19 individuals sequenced deeply classifying ICU patients as healthy (Figure 6b). Curious about the reasons for this, we investigated feature importance and discovered that the two taxonomic parameters “GMHI_good” and “GMHI_bad” both had negative values. This was indicative of the detrimental influence of taxonomy-based features on the performance of the model. As suspected, retraining the Q2PD without them led to the expected, correct predictions (Figure 6c) suggesting that this dataset could be classified based on functional information alone. We further validated this when we performed classification using the taxonomy-based GMHI (Figure 6d) and the hiPCA (Figure 7e), which was inaccurate for GMHI and opposite for hiPCA.

**Figure 7.**
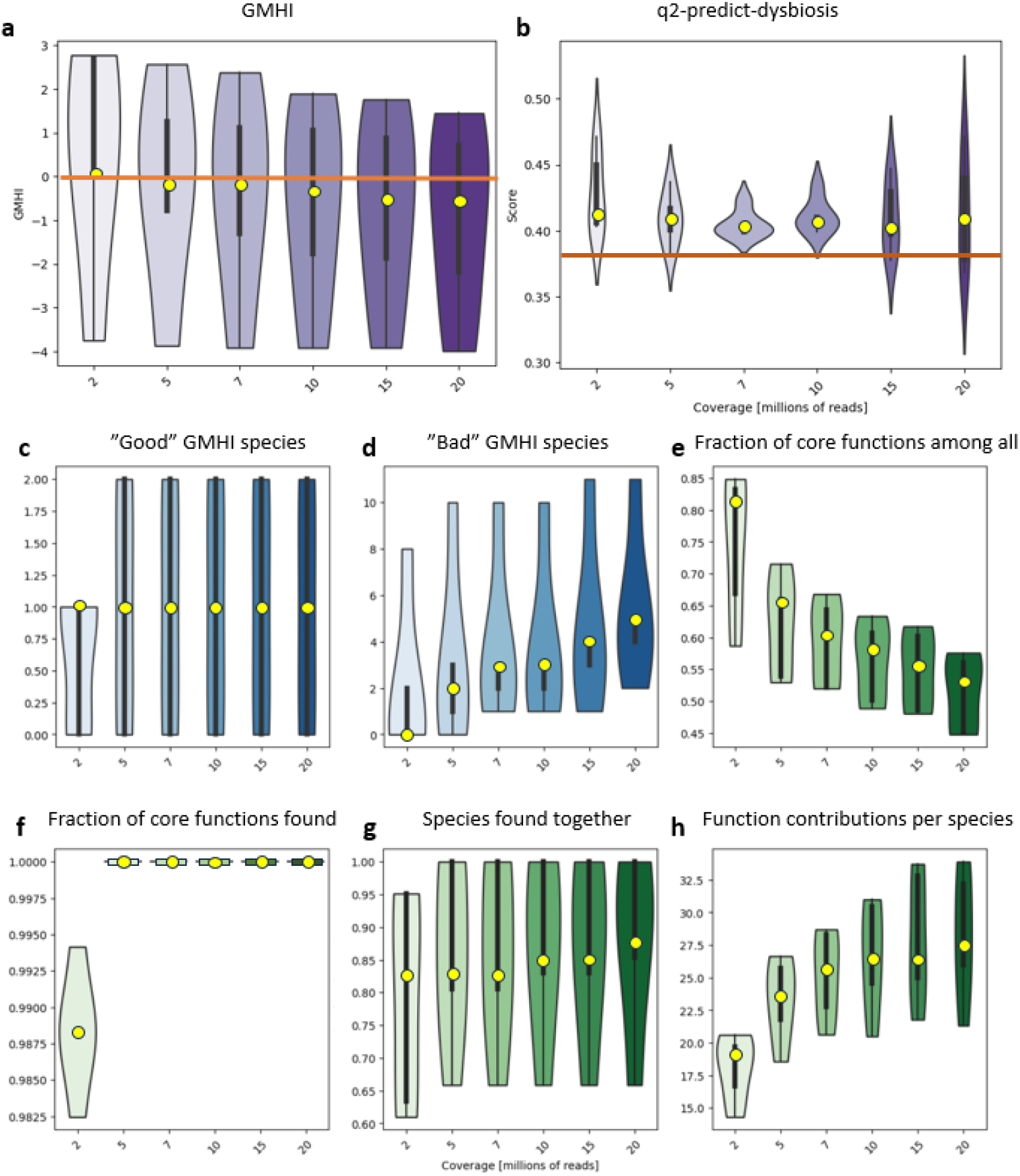
Robustness of health predictions to sequencing coverage. `GMHI` **(a)** and `Q2PD` **(b)** scores for deeply sequenced healthy samples from the COVID-19 cohort, rarefied to corresponding depths. The horizontal lines represent health thresholds relevant to the corresponding methods. Taxonomic `GMHI`-inspired `Q2PD` features **(c, d)** remain relatively stable above the coverage of 5 million reads. Functional and species interaction-related features are much more sensitive **(e-h)**.

Q2PD performed poorly on the shallow COVID-19 dataset which led us to ask where the threshold lay for sequencing depth reliability. In order to investigate method robustness to depth, we applied various degrees of rarefaction to deeply sequenced control samples expecting to see similar prediction scores despite variable sequencing depth. Because the performance of the `hiPCA` (second best classifier after the Q2PD overall) on the COVID-19 cohort was worse than that of the `GMHI`, we decided to use the latter as our benchmark.

The scores produced by the Q2PD were robust and consistent regardless of sequencing depth, whereas for `GMHI`, they increased with degree of rarefaction(Figure 7a,b). Furthermore, the `GMHI` classification appeared to be only relevant for a foreboding depth of 2 million reads (score > 0), as the healthy samples were defined as unhealthy at greater sequencing depth (score < 0). Our index correctly identified healthy samples at any sequencing depth (scores were always above threshold). We plotted the values for each feature separately and found that at low sequencing depths certain species (Figure 7 c, d) or “core functions” (Figure 87e, f) were undetectable. The “Fraction of core features among other features” and the “Function contributions per species” were those most dependent on coverage, which was anticipated due to their dependence on low-abundance functions (Figure 7 e - h). Interestingly, the numbers of “good” and “bad” species identified at any sequencing depth covered only 32% of the complete `GMHI` list (2 and 14, versus 7 and 43 for good and bad species, respectively). In line with the underlying theme of our work and in agreement with all other presented data, this overlap argues strongly against indices that are based solely on taxonomy.

## Discussion

A connection between the human gut microbiome and gut health is now well established, and a number of approaches have been employed in an effort to identify gut dysbiosis from sequence-based analysis of stool samples. Those methods are based on taxonomy and rely either on measures of microbiome richness (alpha / beta diversity) or on the presence or absence of so-called “good” and “bad” bacteria, with the health-indexes `GMHI` (Gupta et al., 2020) and `hiPCA` (Zhu et al., 2023) proposed formally. Hampering such approaches, however, are ecological considerations of metabolic or functional redundancies inherent within complex environments. Currently, re-defining the microbiome to include **inferred functionality** is bolstered by recent studies that highlight the importance of interactions between microbiome components and functional aspects thereof. To address the inadequacy of dogmatic *Linnéan* (taxonomy-based) approach for evaluating change in microbiomes, we developed a novel method that incorporates function in bioinformatics-based assessment of microbiome dysbioses. We show that features based on microbiome functions and interactions define a healthy microbiome more accurately. There exists a set of “core functions”, which are consistently identified as present in healthy gut microbiomes, and which disappear in the advent of dysbiosis. We compare our results to the **hiPCA**, the `GMHI` and two `Shannon entropy` measures (calculated on either species or on functions). Our index not only outperforms the other methods in the originally targeted IBD classification, but also when applied to a range of other diseases. Finally, it is robust to sequencing depth, unlike the `GMHI`, despite being based on sequencing depth-sensitive parameters.

While we here present the index `Q2PD`, we acknowledge that challenges remain prior to eventual clinical deployment as the method’s robustness for diseases other than IBD and obesity is improved. One of the most fascinating results from this work, apart from the model itself, was the finding that different parameters were of varying importance across different diseases and cohorts. This finding could have practical implications in the clinic, as it could indicate the directionality of the microbiome-disease connections. By this we mean specifically that diseases originating in the gut, e.g. IBD, are usually associated with taxonomic shifts and thus better classified with taxonomy-based indices while diseases originating elsewhere may have (functional) effects on the gut and thus better identified with function-oriented methods. It should be noted though that the etiology of many diseases, i.e. whether they originate in the gut or not, remains unknown.

Clearly, a deeper understanding of how the identified function- and interaction-based microbiome features respond to variations in sequencing depth and quality is required. Our initial test did find that longitudinal data provides more insight into personalized “core features” of a microbiome, and this may be indicative of individual deviations from the normal. Thus, tracking microbiome changes in individuals over time may facilitate the early identification of microbiome trajectories or alterations that could help determine risk for certain conditions or diseases, potentiating personalized clinical intervention.

A caveat to this type of study is the infancy of certain fields, in particular, the availability of functional annotation software. Currently, the only widely accepted functional annotation software is HUMAnN (any version), which requires data to be processed in a particular way. In future work we plan to explore other possibilities and ultimately migrate to other inputs (such as mifaser) as well as towards a more universal solution, such as gene content (Zhu et al., 2017). Only then can recent developments in augmented functional annotation be efficiently utilized, as we already have shown in other applications (Maranga et al., 2023, Koehler Leman et al., 2023, Szydlowski et al., 2023).

We are also considering the introduction of a multi-omics approach to improve our predictions, combining metagenomics, metabolomics and proteomics data. With decreasing costs for each of the above, predicting patient health status based on all three −omics methods might be a reasonable option that improves accuracy and prediction in the not-too-distant future. It would also allow us to quantitatively estimate the functions directly, instead of basing our analyses on the functional potential evaluated based on metagenomics data. As a primer for our current efforts, the integration of multiple levels could be performed using the aforementioned MDFS, by investigating relationships between the layers, or using more advanced network-based approaches such has ViLoN (Kańduła et al., 2023).

In conclusion, we highlight the performance and high accuracy obtained by **Q2PD**, based only on a limited number of highly relevant parameters. Despite its development to separate samples derived from healthy vs IBD individuals, its broad applicability was proven to accurately distinguish between healthy patients and individuals with other disease states and conditions. Motivated by this success we are convinced that our approach and methodology provide a better mechanistic understanding of the human gut microbiome and its health, one based on resource competition and interactions based on fundamental ecological principles that ultimately define microbial community structure.

## Methods

### Defining metagenomic parameters associated with health

The raw shotgun sequencing fastq files for the HMP2 and the two IBD validation cohorts used to derive metagenomic health-associated parameters were processed with **Trim Galore 0**.**6**.**10** (Krueger, 2023) to ensure sufficient read quality. The taxonomic profiles were calculated with **MetaPhlAn 4**.**0**.**6** (Blanco-Míguez et al., 2023) and functional profiles with **HUMAnN 3**.**7** (Beghini et al., 2021). Species selection was performed using MultiDimentional Feature Selection, or the `MDFS` (Piliszek et al., 2019, Mnich & Rudnicki, 2020) with a Benjamini-Hochberg p-value correction in a 2D mode. Selection was performed per cohort (healthy versus each of the non-healthy groups in separate `MDFS` runs), and the final list of species was the union of the MDFS results for all cohorts (corrected p-values < 0.05). Metagenomic table formatting and alpha diversity plots were done using `QIIME 2` (Bolyen et al., 2019) and `SparCC` correlations with the `SCNIC` plugin (https://github.com/lozuponelab/q2-SCNIC). Plots were made using custom python scripts. Any statistical tests were calculated with the independent t-test, unless stated otherwise.

### IBD index

Random Forest is a classification and regression method (Breiman, 2001). The algorithm utilises an ensemble of CART trees (Leo Breiman, 1984), in which each tree is built using different bootstrap samples of data and different random subsets of variables at each stage of the tree construction. It is a robust and versatile algorithm that works well on different types of data (Fernandez-Delgado et al., 2014).

Metagenomic features passed to the random forest were constructed based on healthy samples from the HMP2 and two validation cohorts (Val_1 and Val_2). The features are presented in Table 1.

The importance of the features was assessed using a permutation approach and 5-fold cross validation.

From this step on, we used the manually curated samples from the curatedMetagenomicsRepository to ensure consistent sample processing (MetaPhlAn (Blanco-Míguez et al., 2023) and HUMAnN (Beghini et al., 2021), specifically we used the MetaCyc functional annotations) as well as to provide an additional layer of validation of our approach. Our manual curation excluded a samples for the following reasons: incomplete metadata, use of antibiotics or other similar drugs, repeats from the same patients, young age (newborns), or healthy samples not being fully healthy (high BMI, chronic conditions). During filtering, we set the following thresholds: species abundance >= 0.1%, pathway coverage >= 20% and function abundance >= 0.01% - with any features below that changed to 0.

During random forest training, healthy individuals were labeled as “1” and the IBD as “0”. A leave-one-out cross-validation procedure was applied to predict health scores for each sample in order to avoid model overfitting or the need to split the data into training and testing sets. The final health score is the output of the random forest’s *predict_proba* method, which expresses the probability of each sample belonging to the healthy (or unhealthy) group.

The GMHI scores were calculated with the `QIIME 2 q2-health-index` plugin (“q2-health-index,” 2023) and the `hiPCA` (Zhu et al., 2023) predictions were obtained by substituting the original test file with our test samples, due to a lack of instructions about how to do it better (this could potentially result in an overlap of the train and test samples, possibly falsely increasing the `hiPCA`’s accuracy). The Shannon entropies were calculated separately on taxonomic and functional profiles for each sample using custom python scripts.

The GMHI was the only parameter with pre-defined threshold for health. In the case of the other methods, the optimal threshold would be defined using the Youden’s J statistic calculated on the training data.

The relevance of each index in the context of IBD predictions was calculated using `Boruta` (Kursa & Rudnicki, 2010). The Boruta algorithm is a wrapper around the Random Forest classifier. It works in the following way: a data set is extended by adding a randomly permuted copy of each original variable - a so-called shadow variable. A predictive model for the decision variable is then built using the random forest algorithm. The importance of each variable is estimated using a permutation test. Boruta collects the information about the importance and then compares the importance score of each original variable with the maximal importance achieved by a shadow variable.

The procedure is repeated multiple times, and a statistical test is performed. The variables are eventually assigned to three classes: Confirmed (better than random), Rejected (no better than random) and Tentative (those that could not be assigned neither to Confirmed no to the Tentative class. The utility of the variable is a good indicator of the information importance carried by the variable, also in situations where the synergistic interactions are important.

In the Boruta section of our analysis, the predictions of each index were passed as parameters and the health labels (0/1) were used as decision vectors. Index importances were redefined as ranks – with the highest importance marked as rank “1”.

### Index testing: expanding to other diseases

For the purpose of testing, the index was re-trained on the complete set of 30 datasets. However, no parameters were modified. To ensure a reliable benchmark, a leave-one-cohort-out procedure with a 5-fold cross validation was performed. The accuracy and AUC statistics were defined separately for each test cohort based on the validation set (which was a subset of the training dataset).

The performance of our index was evaluated by calculating its winning margin over hiPCA, the second best index, and the mean of the other methods. Specifically, for each test dataset, the AUC of the hiPCA or the mean AUC of the other methods was subtracted from the AUC produced by the Q2PD. The winning margin was defined as a mean of the AUC differences, separately for cases when Q2PD won and when not.

Importance of the features contributing to the Q2PD score was evaluated by training a random forest model separately for each cohort and using a feature permutation approach. The importances were changed to ranks, with the highest ranks (lowest values) representing the greatest importance.

### Final index and application to the COVID-19 dataset

The final index was trained on the complete set of 30 cohorts using 5-fold cross validation. The health threshold was defined based on a mean of thresholds defined on training data for each iteration of the leave-one-cohort-out approach and was set at 0.38 (which was very similar for all iterations).

To evaluate the indices in the context of the robustness to sequencing depth, healthy deeply sequenced samples from the COVID-19 dataset were rarefied to different sequencing depths using **seqtk 1**.**4** (Li, 2023).

## Data and code availability

The HMP2 data can be found in Sequence Read Archive (SRA) with the accession code PRJNA398089. The two IBD validation cohorts can be located under accessions PRJNA389280 and PRJEB1220. The test cohorts were downloaded from the curatedMetagenomicsData repository (Pasolli et al., 2017). The samples from this repository were manually selected - we removed samples from children, reported control yet unhealthy samples, as well as those of patients undergoing antibiotic treatment. The COVID-19 cohort can be found under the accession PRJEB64515 in the European Nucleotide Archive. The Q2PD is deposited on GitHub and can be accessed along with sample accessions and scripts needed to reproduce our results here: https://github.com/Kizielins/q2-predict-dysbiosis. The biotools identifier of the software is biotools:q2-predict-dysbiosis and the RRID is SCR_026038.

## Acknowledgements

This research was conducted as a part of the NCN Sonata BIS grant number 2020/38/E/NZ2/00598. We gratefully acknowledge Poland’s high-performance Infrastructure PLGrid (HPC Centers: ACK Cyfronet AGH, PCSS, CI TASK, WCSS) for providing computer facilities. We would like to thank Dr Krzysztof Mnich, Dr Sajad Shahbazi, Dr Balakrishnan Subramanian and Piotr Stomma from the University of Białystok for their expertise in the Multi-Dimensional Feature Selection algorithm and insightful contributions to our work, dr. Tomasz Kościółek from the Sano Centre for Computational Medicine and the members of Bioinformatics Research Group at MCB JU for their comments and support.

## Ethics declarations

All authors declare no competing interests.

