## Supplementary materials for "Healthy microbiome - moving towards functional interpretation"

### Supplement

#### Supplementary Table 1

“Core functions”: functions present in at least 80% of healthy individuals from the HMP2 and two validation cohorts

**Core function name**

1CMET2-PWY: folate transformations III (E. coli)

ANAEROFRUCAT-PWY: homolactic fermentation

ANAGLYCOLYSIS-PWY: glycolysis III (from glucose)

ARGININE-SYN4-PWY: L-ornithine biosynthesis II

ARGSYN-PWY: L-arginine biosynthesis I (via L-ornithine)

ARGSYNBSUB-PWY: L-arginine biosynthesis II (acetyl cycle)

ARO-PWY: chorismate biosynthesis I

ASPASN-PWY: superpathway of L-aspartate and L-asparagine biosynthesis

BIOTIN-BIOSYNTHESIS-PWY: biotin biosynthesis I

BRANCHED-CHAIN-AA-SYN-PWY: superpathway of branched chain amino acid biosynthesis

CALVIN-PWY: Calvin-Benson-Bassham cycle

CITRULBIO-PWY: L-citrulline biosynthesis

COA-PWY-1: superpathway of coenzyme A biosynthesis III (mammals)

COA-PWY: coenzyme A biosynthesis I (prokaryotic)

COBALSYN-PWY: superpathway of adenosylcobalamin salvage from cobinamide I

COLANSYN-PWY: colanic acid building blocks biosynthesis

COMPLETE-ARO-PWY: superpathway of aromatic amino acid biosynthesis

DTDPRHAMSYN-PWY: dTDP-&beta;-L-rhamnose biosynthesis

FASYN-ELONG-PWY: fatty acid elongation -- saturated

FERMENTATION-PWY: mixed acid fermentation

FUCCAT-PWY: fucose degradation

GALACTUROCAT-PWY: D-galacturonate degradation I

GLCMANNANAUT-PWY: superpathway of N-acetylglucosamine, N-acetylmannosamine and N-acetylneuraminate degradation

GLUCONEO-PWY: gluconeogenesis I

GLUCUROCAT-PWY: superpathway of &beta;-D-glucuronosides degradation

GLUTORN-PWY: L-ornithine biosynthesis I

GLYCOCAT-PWY: glycogen degradation I

GLYCOGENSYNTH-PWY: glycogen biosynthesis I (from ADP-D-Glucose)

GLYCOLYSIS-E-D: superpathway of glycolysis and the Entner-Doudoroff pathway

GLYCOLYSIS: glycolysis I (from glucose 6-phosphate)

HISDEG-PWY: L-histidine degradation I

HISTSYN-PWY: L-histidine biosynthesis

HSERMETANA-PWY: L-methionine biosynthesis III

ILEUSYN-PWY: L-isoleucine biosynthesis I (from threonine)

NAGLIPASYN-PWY: lipid IVA biosynthesis (E. coli)

NONMEVIPP-PWY: methylerythritol phosphate pathway I

NONOXIPENT-PWY: pentose phosphate pathway (non-oxidative branch) I

OANTIGEN-PWY: O-antigen building blocks biosynthesis (E. coli)

P41-PWY: pyruvate fermentation to acetate and (S)-lactate I

PANTO-PWY: phosphopantothenate biosynthesis I

PANTOSYN-PWY: superpathway of coenzyme A biosynthesis I (bacteria)

PENTOSE-P-PWY: pentose phosphate pathway

PEPTIDOGLYCANSYN-PWY: peptidoglycan biosynthesis I (meso-diaminopimelate containing)

PHOSLIPSYN-PWY: superpathway of phospholipid biosynthesis I (bacteria)

POLYISOPRENSYN-PWY: polyisoprenoid biosynthesis (E. coli)

PWY-1042: glycolysis IV

PWY-1269: CMP-3-deoxy-D-manno-octulosonate biosynthesis

PWY-2941: L-lysine biosynthesis II

PWY-2942: L-lysine biosynthesis III

PWY-3001: superpathway of L-isoleucine biosynthesis I

PWY-3841: folate transformations II (plants)

PWY-4984: urea cycle

PWY-5030: L-histidine degradation III

PWY-5097: L-lysine biosynthesis VI

PWY-5100: pyruvate fermentation to acetate and lactate II

PWY-5103: L-isoleucine biosynthesis III

PWY-5121: superpathway of geranylgeranyl diphosphate biosynthesis II (via MEP)

PWY-5130: 2-oxobutanoate degradation I

PWY-5154: L-arginine biosynthesis III (via N-acetyl-L-citrulline)

PWY-5188: tetrapyrrole biosynthesis I (from glutamate)

PWY-5384: sucrose degradation IV (sucrose phosphorylase)

PWY-5484: glycolysis II (from fructose 6-phosphate)

PWY-5505: L-glutamate and L-glutamine biosynthesis

PWY-5659: GDP-mannose biosynthesis

PWY-5667: CDP-diacylglycerol biosynthesis I

PWY-5686: UMP biosynthesis I

PWY-5695: inosine 5'-phosphate degradation

PWY-5941: glycogen degradation II

PWY-5973: cis-vaccenate biosynthesis

PWY-5989: stearate biosynthesis II (bacteria and plants)

PWY-6121: 5-aminoimidazole ribonucleotide biosynthesis I

PWY-6122: 5-aminoimidazole ribonucleotide biosynthesis II

PWY-6123: inosine-5'-phosphate biosynthesis I

PWY-6124: inosine-5'-phosphate biosynthesis II

PWY-6125: superpathway of guanosine nucleotides de novo biosynthesis II

PWY-6126: superpathway of adenosine nucleotides de novo biosynthesis II

PWY-6147: 6-hydroxymethyl-dihydropterin diphosphate biosynthesis I

PWY-6151: S-adenosyl-L-methionine salvage I

PWY-6163: chorismate biosynthesis from 3-dehydroquinate

PWY-621: sucrose degradation III (sucrose invertase)

PWY-6270: isoprene biosynthesis I

PWY-6277: superpathway of 5-aminoimidazole ribonucleotide biosynthesis

PWY-6282: palmitoleate biosynthesis I (from (5Z)-dodec-5-enoate)

PWY-6292: superpathway of L-cysteine biosynthesis (mammalian)

PWY-6305: superpathway of putrescine biosynthesis

PWY-6317: D-galactose degradation I (Leloir pathway)

PWY-6353: purine nucleotides degradation II (aerobic)

PWY-6385: peptidoglycan biosynthesis III (mycobacteria)

PWY-6386: UDP-N-acetylmuramoyl-pentapeptide biosynthesis II (lysine-containing)

PWY-6387: UDP-N-acetylmuramoyl-pentapeptide biosynthesis I (meso-diaminopimelate containing)

PWY-6507: 4-deoxy-L-threo-hex-4-enopyranuronate degradation

PWY-6519: 8-amino-7-oxononanoate biosynthesis I

PWY-6527: stachyose degradation

PWY-6606: guanosine nucleotides degradation II

PWY-6608: guanosine nucleotides degradation III

PWY-6609: adenine and adenosine salvage III

PWY-6628: superpathway of L-phenylalanine biosynthesis

PWY-6630: superpathway of L-tyrosine biosynthesis

PWY-6700: queuosine biosynthesis I (de novo)

PWY-6703: preQ0 biosynthesis

PWY-6731: starch degradation III

PWY-6823: molybdopterin biosynthesis

PWY-6859: all-trans-farnesol biosynthesis

PWY-6897: thiamine diphosphate salvage II

PWY-6901: superpathway of glucose and xylose degradation

PWY-6902: chitin degradation II (Vibrio)

PWY-6936: seleno-amino acid biosynthesis (plants)

PWY-6969: TCA cycle V (2-oxoglutarate synthase)

PWY-702: L-methionine biosynthesis II

PWY-7111: pyruvate fermentation to isobutanol (engineered)

PWY-7197: pyrimidine deoxyribonucleotide phosphorylation

PWY-7199: pyrimidine deoxyribonucleosides salvage

PWY-7208: superpathway of pyrimidine nucleobases salvage

PWY-7220: adenosine deoxyribonucleotides de novo biosynthesis II

PWY-7221: guanosine ribonucleotides de novo biosynthesis

PWY-7222: guanosine deoxyribonucleotides de novo biosynthesis II

PWY-7228: superpathway of guanosine nucleotides de novo biosynthesis I

PWY-7229: superpathway of adenosine nucleotides de novo biosynthesis I

PWY-7234: inosine-5'-phosphate biosynthesis III

PWY-7237: myo-, chiro- and scyllo-inositol degradation

PWY-7238: sucrose biosynthesis II

PWY-7242: D-fructuronate degradation

PWY-724: superpathway of L-lysine, L-threonine and L-methionine biosynthesis II

PWY-7282: 4-amino-2-methyl-5-diphosphomethylpyrimidine biosynthesis II

PWY-7323: superpathway of GDP-mannose-derived O-antigen building blocks biosynthesis

PWY-7328: superpathway of UDP-glucose-derived O-antigen building blocks biosynthesis

PWY-7345: superpathway of anaerobic sucrose degradation

PWY-7357: thiamine phosphate formation from pyrithiamine and oxythiamine (yeast)

PWY-7383: anaerobic energy metabolism (invertebrates, cytosol)

PWY-7392: taxadiene biosynthesis (engineered)

PWY-7400: L-arginine biosynthesis IV (archaebacteria)

PWY-7456: &beta;-(1,4)-mannan degradation

PWY-7560: methylerythritol phosphate pathway II

PWY-7663: gondoate biosynthesis (anaerobic)

PWY-7664: oleate biosynthesis IV (anaerobic)

PWY-7761: NAD salvage pathway II (PNC IV cycle)

PWY-7790: UMP biosynthesis II

PWY-7791: UMP biosynthesis III

PWY-7851: coenzyme A biosynthesis II (eukaryotic)

PWY-7953: UDP-N-acetylmuramoyl-pentapeptide biosynthesis III (meso-diaminopimelate containing)

PWY-7977: L-methionine biosynthesis IV

PWY-8004: Entner-Doudoroff pathway I

PWY-8073: lipid IVA biosynthesis (P. putida)

PWY-8178: pentose phosphate pathway (non-oxidative branch) II

PWY-8187: L-arginine degradation XIII (reductive Stickland reaction)

PWY-841: superpathway of purine nucleotides de novo biosynthesis I

PWY-I9: L-cysteine biosynthesis VI (from L-methionine)

PWY0-1241: ADP-L-glycero-&beta;-D-manno-heptose biosynthesis

PWY0-1261: anhydromuropeptides recycling I

PWY0-1296: purine ribonucleosides degradation

PWY0-1319: CDP-diacylglycerol biosynthesis II

PWY0-1586: peptidoglycan maturation (meso-diaminopimelate containing)

PWY0-162: superpathway of pyrimidine ribonucleotides de novo biosynthesis

PWY0-845: superpathway of pyridoxal 5'-phosphate biosynthesis and salvage

PWY0-862: (5Z)-dodecenoate biosynthesis I

PWY4FS-7: phosphatidylglycerol biosynthesis I (plastidic)

PWY4FS-8: phosphatidylglycerol biosynthesis II (non-plastidic)

PWY66-399: gluconeogenesis III

PWY66-429: fatty acid biosynthesis initiation (mitochondria)

PYRIDNUCSYN-PWY: NAD de novo biosynthesis I (from aspartate)

PYRIDOXSYN-PWY: pyridoxal 5'-phosphate biosynthesis I

RHAMCAT-PWY: L-rhamnose degradation I

RIBOSYN2-PWY: flavin biosynthesis I (bacteria and plants)

SALVADEHYPOX-PWY: adenosine nucleotides degradation II

SER-GLYSYN-PWY: superpathway of L-serine and glycine biosynthesis I

THISYNARA-PWY: superpathway of thiamine diphosphate biosynthesis III (eukaryotes)

THRESYN-PWY: superpathway of L-threonine biosynthesis

TRNA-CHARGING-PWY: tRNA charging

UDPNAGSYN-PWY: UDP-N-acetyl-D-glucosamine biosynthesis I

VALSYN-PWY: L-valine biosynthesis

#### Supplementary Figure 1

Number of observed functions per sample, separated by health group and project.

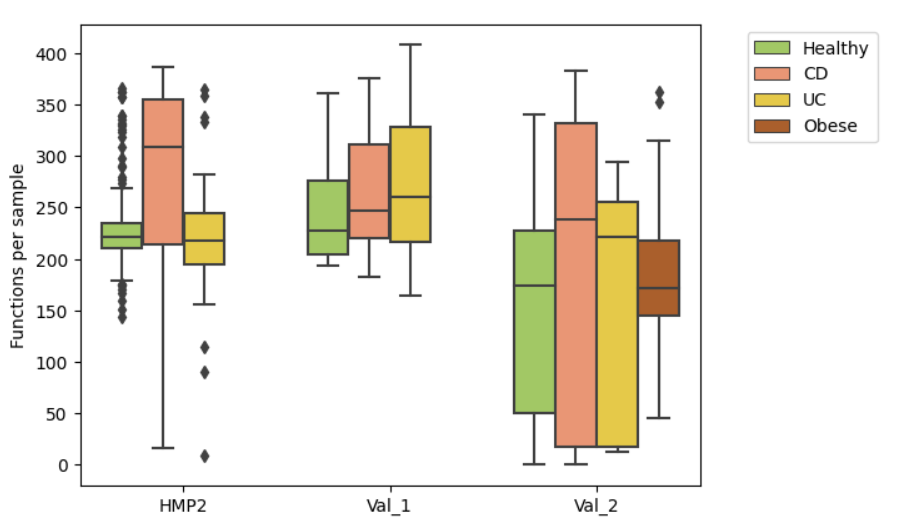

#### Supplementary Figure 2

Comparison of core functions in health and disease. **a.** Overlap of LEfSe differentially enriched pathways in healthy versus unhealthy samples in the validation and HMP2 cohorts. **b.** Overlap of LEfSe differentially enriched pathways in unhealthy versus healthy samples in the validation and HMP2 cohorts**. c.** Fraction of seen core functions among all annotated core functions. **d.** Fraction of seen core functions among all functions seen in a sample.

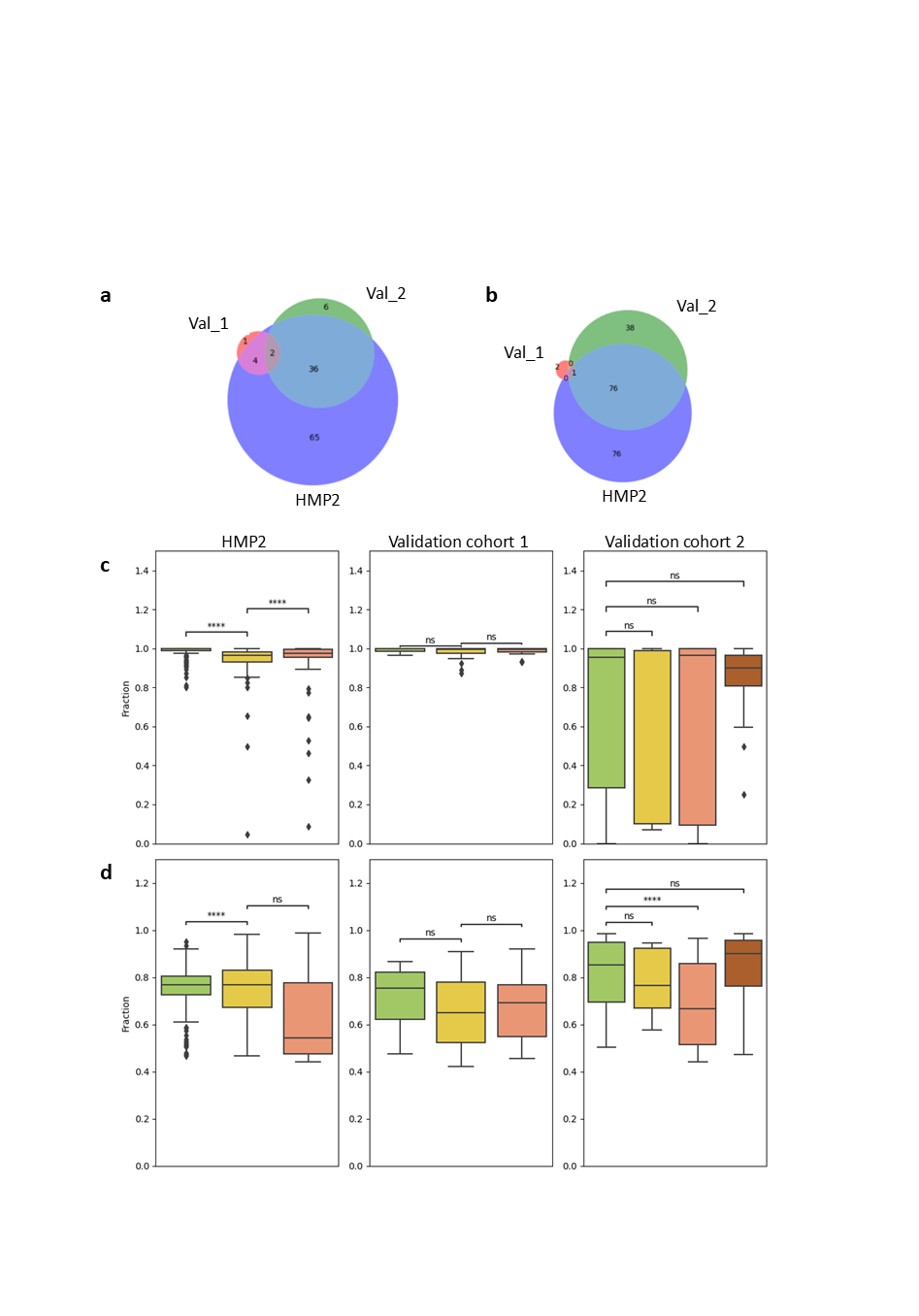

#### Supplementary Figure 3

Number of observed species per sample, separated by health group and project.

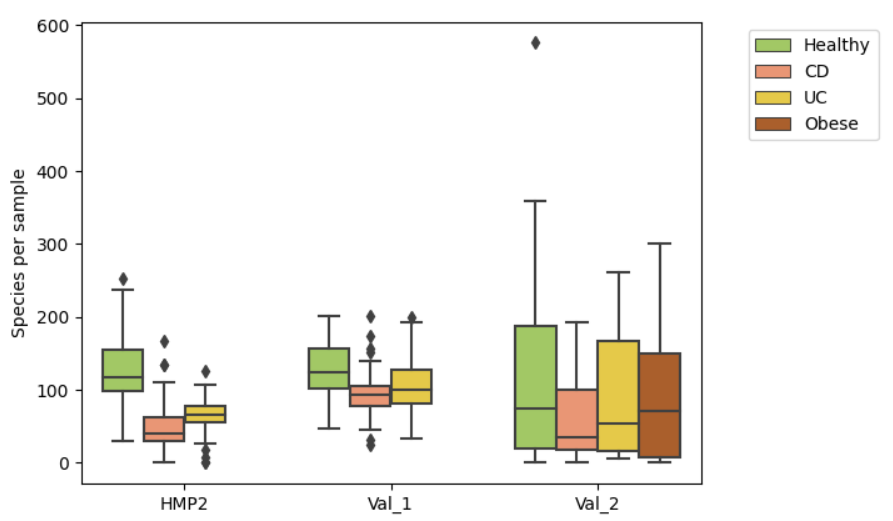

#### Supplementary Figure 4

Top species contributions to core functions, based on the stratified output of HUMAnN. For every cohort, the sum of the contributions for all samples was calculated and contributions with the greatest summed abundances are shown. The final values have been normalized to values between 0 and 1 to compare the strength of the correlations between the groups.

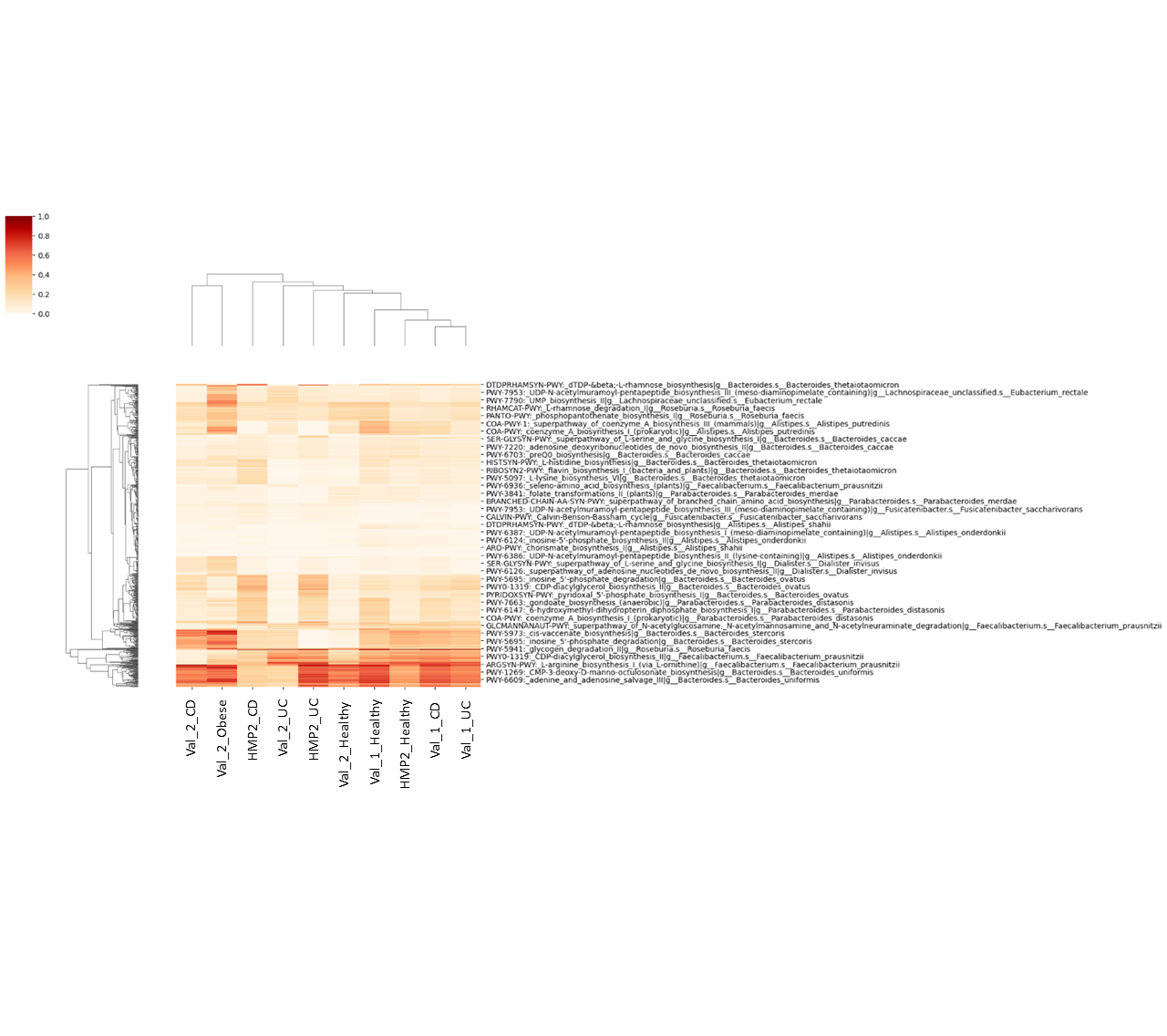

#### Supplementary Figure 5

Functional redundancy. **a.** Average number of species per function in a sample. **b**. Average number of functions per species.

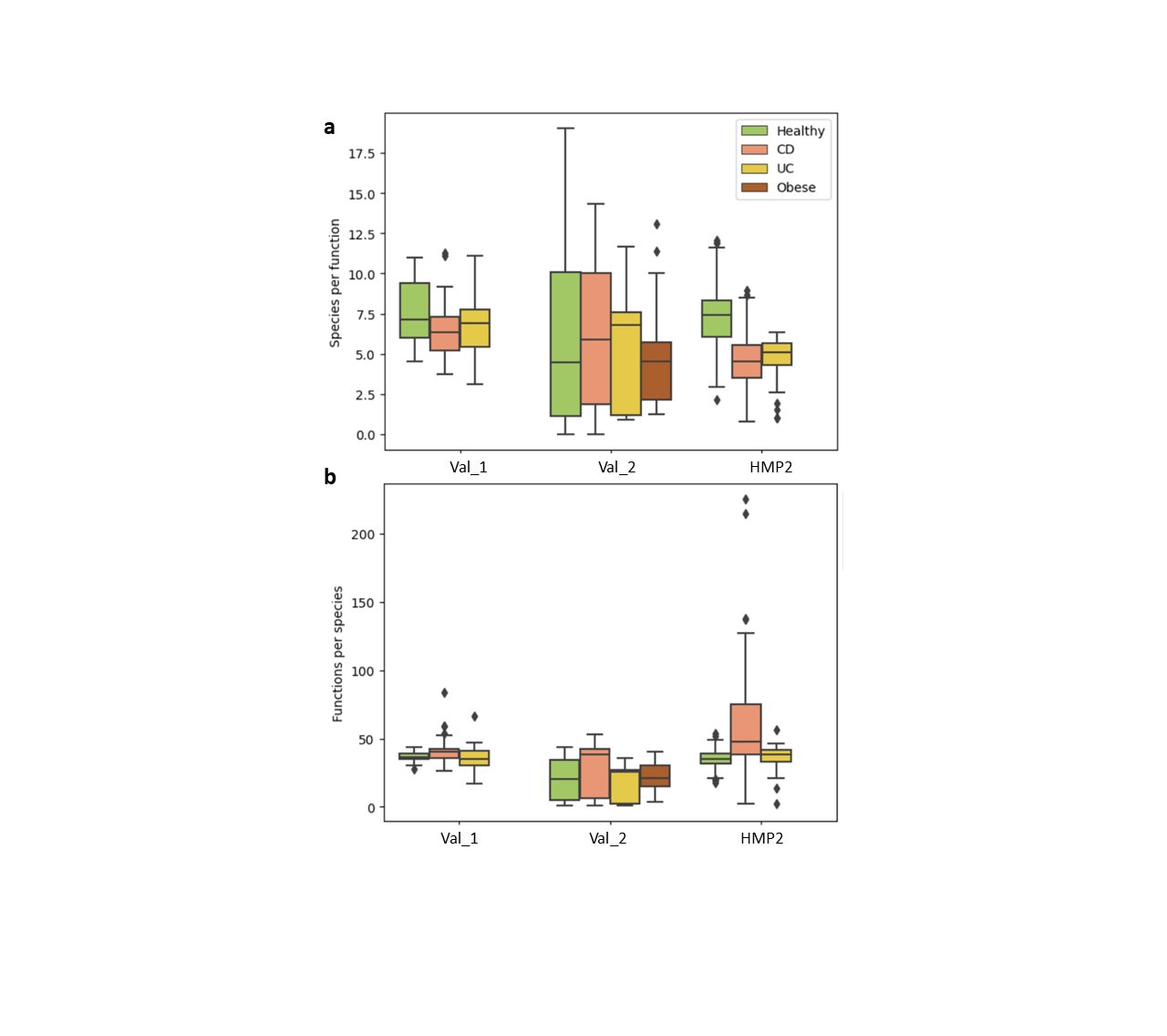

#### Supplementary Figure 6

Accuracy and AUC scores achieved by each index, separately for each cohort.

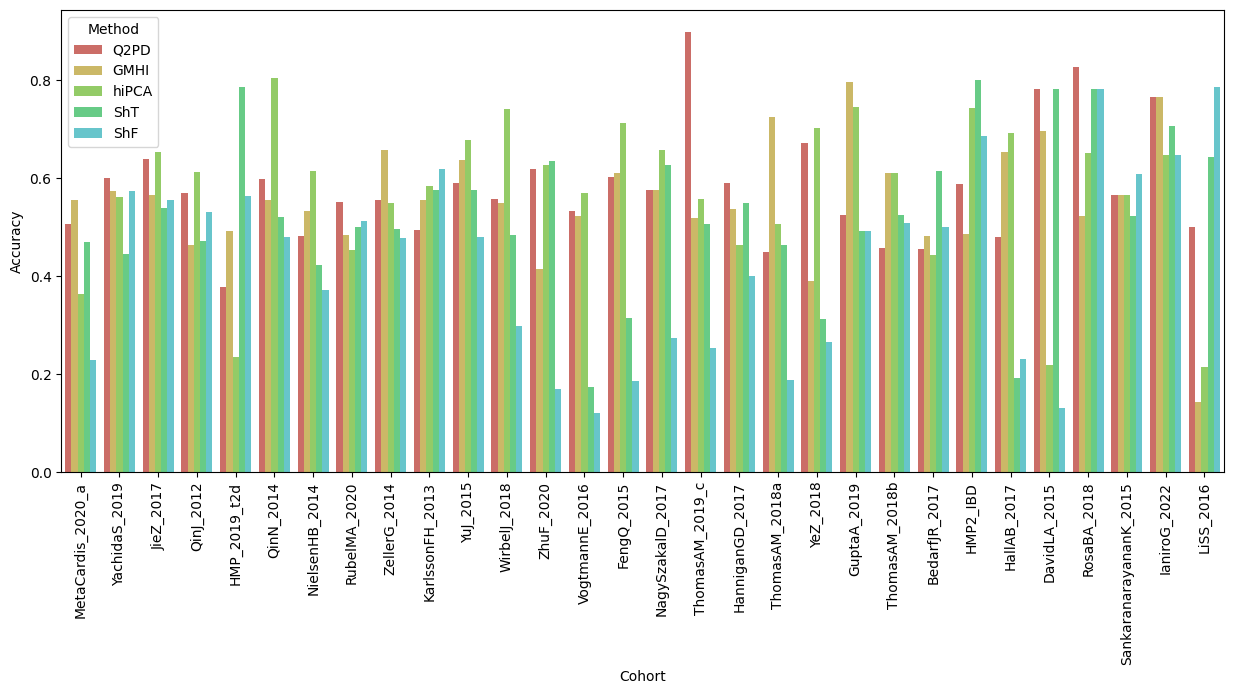

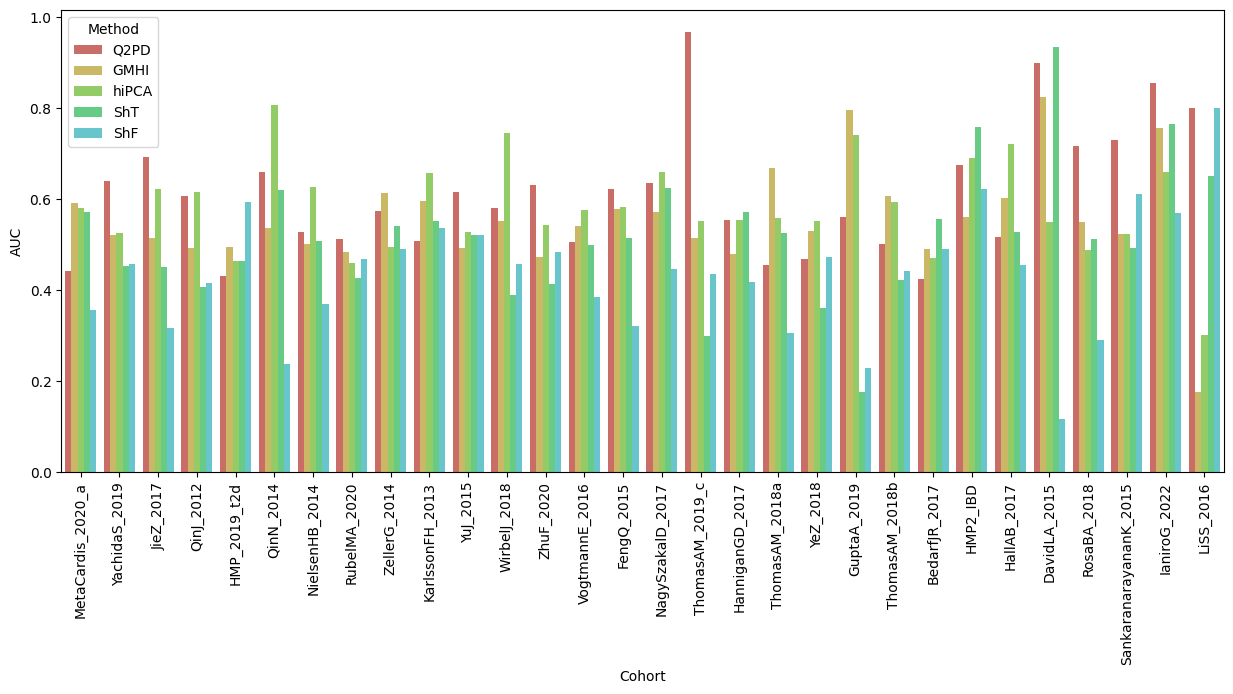
